# Inhibition mediated by group III mGluRs regulates habenula activity and defensive behaviors

**DOI:** 10.1101/2024.09.11.612421

**Authors:** Anna Maria Ostenrath, Nicholas Faturos, Yağnur Işık Çiftci Çobanoğlu, Bram Serneels, Inyoung Jeong, Anja Enz, Francisca Hinrichsen, Aytac Kadir Mutlu, Ricarda Bardenhewer, Suresh Kumar Jetti, Stephan C. F. Neuhauss, Nathalie Jurisch-Yaksi, Emre Yaksi

**Affiliations:** Kavli Institute for Systems Neuroscience and Center for Algorithms in the Cortex, Norwegian University of Science and Technology, Trondheim, Norway; Koç University Research Center for Translational Medicine, Koç University School of Medicine, Istanbul, Turkey; University of Zurich, Department of Molecular Life Sciences, Zurich, Switzerland; Department of Clinical and Molecular Medicine, Norwegian University of Science and Technology, Trondheim, Norway; Neuro-Electronics Research Flanders, Leuven, Belgium; Medical Scientist Training Program, University of Minnesota School of Medicine, Minneapolis, MN, 55455, USA

## Abstract

Inhibition contributes to various brain computations from sensory motor transformations to cognitive operations. While most studies on inhibition focus on GABA, the main excitatory neurotransmitter of the brain, glutamate, can also elicit inhibition via metabotropic glutamate receptors (mGluRs). The function of mGluR-mediated inhibition remains largely elusive. Here, we investigated the role of group III mGluR-dependent inhibition in the habenula. This primarily glutamatergic and conserved forebrain region acts as a hub between multiple forebrain inputs and neuromodulatory mid- and hindbrain targets that regulate adaptive behaviors. We showed that both zebrafish and mice habenula express group III mGluRs. We identified that group III mGluRs regulate the membrane potential and calcium activity of zebrafish dorsal habenula. Pharmacological and genetic perturbation of group III mGluRs increased sensory-evoked excitation and reduced selectivity of habenular neurons to different sensory modalities. We also observed that inhibition is the main channel of communication between primarily glutamatergic habenula neurons. Blocking group III mGluRs reduced inhibition within habenula and increased correlations during spontaneous activity. In line with such inhibition within habenula, we identified that multi-sensory information is integrated mainly through competition and suppression across habenular neurons, which in part relies on group III mGluRs. Finally, genetic perturbation of a habenula-specific group III mGluR, mGluR6a, amplified neural responses and defensive behaviors evoked by sensory stimulation and environmental changes. Altogether, our results revealed that mGluR driven inhibition is essential in encoding, integration, and communication of information between Hb neurons, ultimately playing a critical role in regulating defensive and adaptive behaviors.

## INTRODUCTION

Inhibition is a crucial part of brain function (*1*). Apart from controlling brain excitability (*2-6*) and stabilizing network activity (*7, 8*) inhibition also contributes to multiple neural computations from refining sensory representations (*9-26*) to mediating high order cognitive processes (*27-33*). Notably, studies in humans using functional brain imaging have highlighted a decrease in brain activity during attention-demanding cognitive tasks globally, except for the area involved in those specific tasks (*34-36*). y-aminobutyric acid (GABA) is the main inhibitory neurotransmitter of the central nervous system (*37*), and is released from diverse classes of GABAergic neurons with various roles in neural computations (*38-40*). Besides GABA, neuromodulators such as dopamine (*41*), serotonin (*42-45*) and acetylcholine (*46, 47*) were also shown to mediate inhibition through their specific inhibitory receptors. Glutamate, which is the main excitatory neurotransmitter of the vertebrate central nervous system (*48, 49*), can also lead to inhibition through the action of group II and group III metabotropic glutamate receptors (mGluRs) (*49-51*). Group II/III mGluRs are expressed throughout the brain (*50, 52-54*), and modulate brain excitability by acting on the pre- and post-synapse of neurons as well as glia (*50, 55, 56*). To date, the role of glutamate-driven inhibition in neural computations and animal behavior remains elusive.

In this study, we investigated the role of glutamatergic inhibition in brain function by specifically focusing on the habenula (Hb) and group III mGluRs. The Hb is a diencephalic nucleus associated with adaptive behaviors (*57-64*) and learning (*58, 61, 63, 65-70*). Hb dysfunction in humans is linked to mood disorders (*71-73*). Hb receives diverse inputs from cortico-limbic (*65, 66, 74-83*) and sensory brain regions (*84-87*), acting as a hub integrating and relaying information to target monoaminergic nuclei that regulate animals behavioral states (*58-60, 62, 69, 88-96*). The evolutionarily conserved (*58, 63, 68, 97*) subdivision of the mammalian medial (MHb) and lateral (LHb) Hb, which correspond to the dorsal (dHb) and ventral Hb (vHb) in zebrafish, mediate distinct functions. The vHb/LHb is involved in motivation (*98, 99*), decision making (*60, 88*), learning (*58, 100*), defensive (*66, 101-103*) and coping behaviors (*57, 92*). The dHb/ MHb is associated with circadian rhythms (*104, 105*), social behaviors (*62*), sensory processing (*84, 104, 106-109*) and fear learning (*59, 63, 68*).

Recent studies have shown that the Hb is composed of molecularly diverse (*110-112*) and spatially organized (*58, 63, 68, 97*) clusters or ensembles (*113*) with distinct developmental origin (*97, 108, 114*) and function (*58, 106-109*). These spatially organized clusters of Hb neurons display correlated spontaneous activity among nearby neurons and robust anti-correlations across Hb clusters, as shown in zebrafish (*106, 108, 109, 115*). Such spatially structured spontaneous Hb activity is largely driven by the cortico-limbic structures of the forebrain (*109*). Besides showing correlated spontaneous activity, neurons in individual Hb clusters also exhibit synchronized responses to odors, light and mechanical stimuli (*106, 108, 109*). Interestingly, odor responses in the Hb were shown to inhibit distinct Hb neurons and dampen the way Hb integrates information from its forebrain inputs (*109*). These sensory-evoked decreases in Hb responses and robust anticorrelations in spontaneous Hb activity suggest that inhibition plays an important role in Hb function. However, the nature and function of inhibition in Hb networks, which is largely a glutamatergic brain region (*112, 116*), is not well understood.

In this study, we showed that sensory stimuli evoke both excitation and inhibition in Hb neurons. We identified the expression of several group III mGluRs, which are inhibitory, in both zebrafish and mouse Hb. We observed that pharmacological blockage and activation of group III mGluRs can increase and decrease membrane potential and calcium signals of zebrafish Hb neurons. Our results revealed prominent inhibitory connections across Hb neurons, which are largely reduced by blocking group III mGluRs. Pharmacological or genetic perturbation of group III mGluRs resulted in increased sensory-evoked excitation and decreased inhibition, leading to reduced selectivity and integration of sensory modalities by Hb neurons. Finally, we identified that genetic perturbation of mGluR6a, which is highly enriched in Hb, led to stronger defensive behaviors and reduced adaptation to environmental changes. Altogether, our results revealed that inhibition via group III mGluRs plays an important role in neural connectivity and sensory computations in the Hb and regulates defensive behaviors.

## RESULTS

### Habenula neurons respond to sensory stimulation with excitation and inhibition

To investigate the extent of sensory-evoked inhibition in Hb, we monitored Hb responses to olfactory, visual and mechanical stimulation, by using volumetric two-photon calcium imaging in 3 weeks old juvenile *Tg(elavl3:GCaMP6s*) zebrafish (*108, 109, 117-119*) expressing GCaMP6s pan-neuronally. We identified that all three sensory modalities evoked both excitation and inhibition in a fraction of neurons, distributed across dorsal and ventral Hb (Figure 1). These results demonstrate that the zebrafish Hb responds to multiple sensory modalities both by excitation and inhibition.

**Figure 1:**
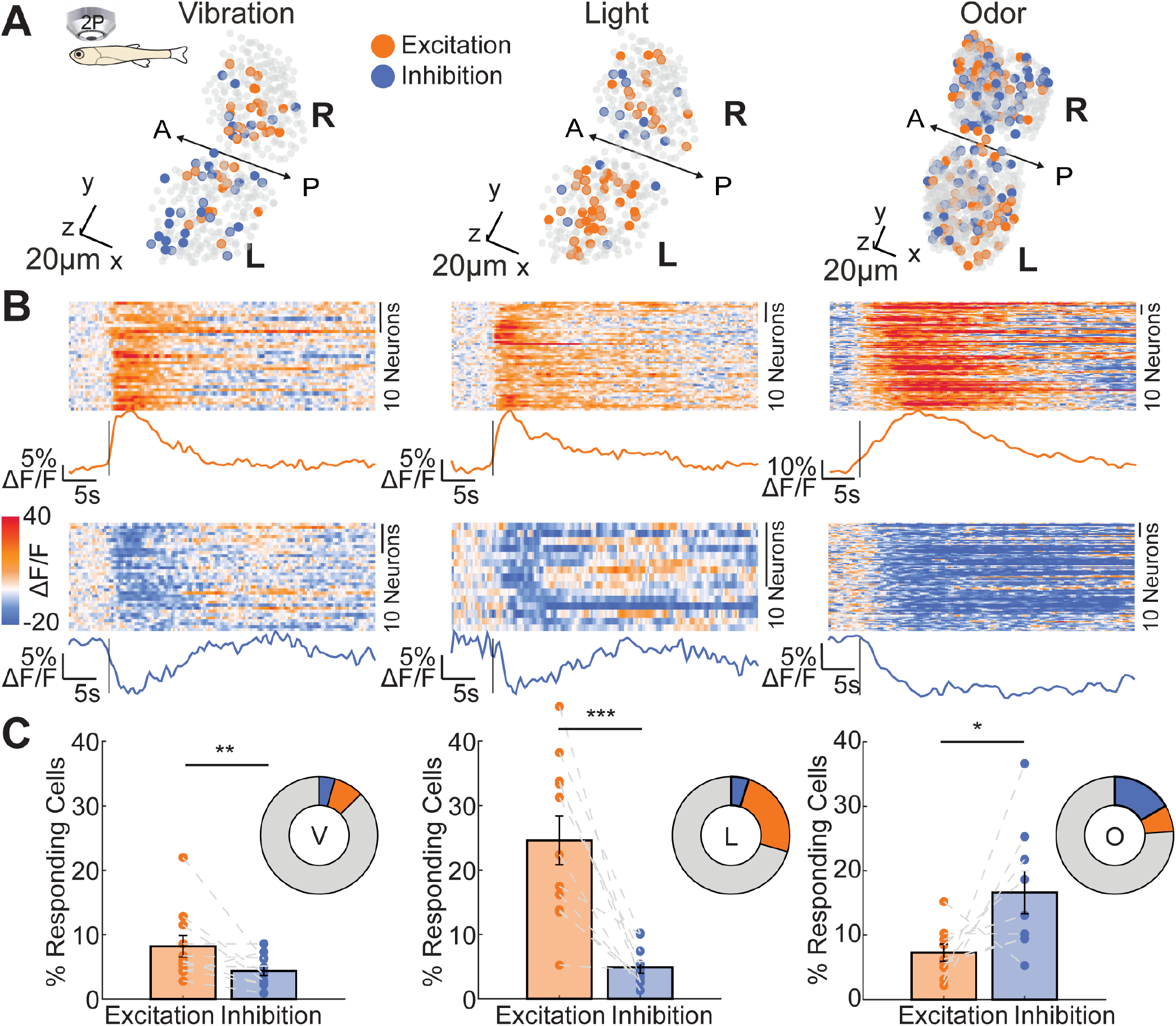
Habenular neurons respond to vibration, light and odors with both excitation and inhibition. (A) Representative examples showing the location of responding neurons in three-dimensional Hb reconstruction in *Tg(elavl3:GCaMP6s*) juvenile zebrafish. Neurons are color coded by their response to vibration (left), light (middle) and odor (amino acid mixture, right). Neurons with activity 2 STD higher than baseline are orange (excitation), 1 STD smaller than baseline are blue (inhibition), non-responding neurons are grey. Scale bar represents 20 μm in x, y and z direction. L, left; R, right; A, anterior; P, posterior. (B) Time-courses of the Hb calcium signals (ΔF/F) of the excited (top) and inhibited (bottom) neurons from panel A to the vibration (left), light (middle) and odor (right) stimulations. Orange and blue color lines represent average excitation and inhibition. (C) Percentage of excited and inhibited cells in Hb upon vibration (left, n=11), light (middle, n=11) and odor (right, n=9) stimulations. (*p < 0.05, ** p < 0.01, *** p < 0.001), tailed Wilcoxon signed-rank test).

### Group III mGluRs are expressed in habenula and mediate inhibition in habenula neurons

A large fraction of vertebrate Hb neurons is glutamatergic (*108, 112, 116*). Moreover, the primary pathways delivering visual (*84, 85*) and olfactory (*86, 120*) information to zebrafish Hb are glutamatergic. Hence, we asked whether inhibitory mGluRs are present in Hb, which could explain part of the sensory-evoked inhibition (Figure 1) and anticorrelations between Hb neurons during spontaneous activity (*106, 109*). We specifically focused on group III mGluR, since a previous study reported diverse expression of these receptors across the zebrafish brain (*121*). To investigate the group III mGluR expression in vertebrate Hb, we examined and compared two recently published single cell RNA sequencing data from zebrafish (*112*) and mice (*116*). We observed prominent expression of mGluR4, mGluR6a, mGluR7 and mGluR8b in zebrafish Hb, with stronger expression in dHb neurons, when compared to vHb (Figure 2A). Similarly, mGluR4 and mGluR7 were expressed prominently in mouse Hb, but with stronger preference to LHb neurons (Figure 2B). In the mouse single cell RNA sequencing data, we could detect only few neurons with mGluR8 expression in LHb, and no expression of mGluR6. Due to low expression levels of GPCRs, detection of mGluR transcripts is difficult with single cell transcriptomic profiling (*122*). Hence, we generated a parallel spatial transcriptome of zebrafish Hb with subcellular spatial resolution (*123-125*). The spatial transcriptomic approach was more effective in detecting mGluR transcripts. Notably, we identified topographically distinct expression of mGluR4, mGluR6a, mGluR8b in the zebrafish dHb (Figure 3C), which is in line with our single cell RNA sequencing results. Horizontal sections across the entire forebrain, revealed that while mGluR4 and mGluR8b are also present in other forebrain regions, mGluR6a expression was relatively specific to zebrafish Hb (Figure 2D), and the olfactory bulbs (Suppl. Figure 1). These results revealed that group III mGluRs are present in both zebrafish and mouse Hb, with different preferences to distinct Hb subregions.

**Figure 2:**
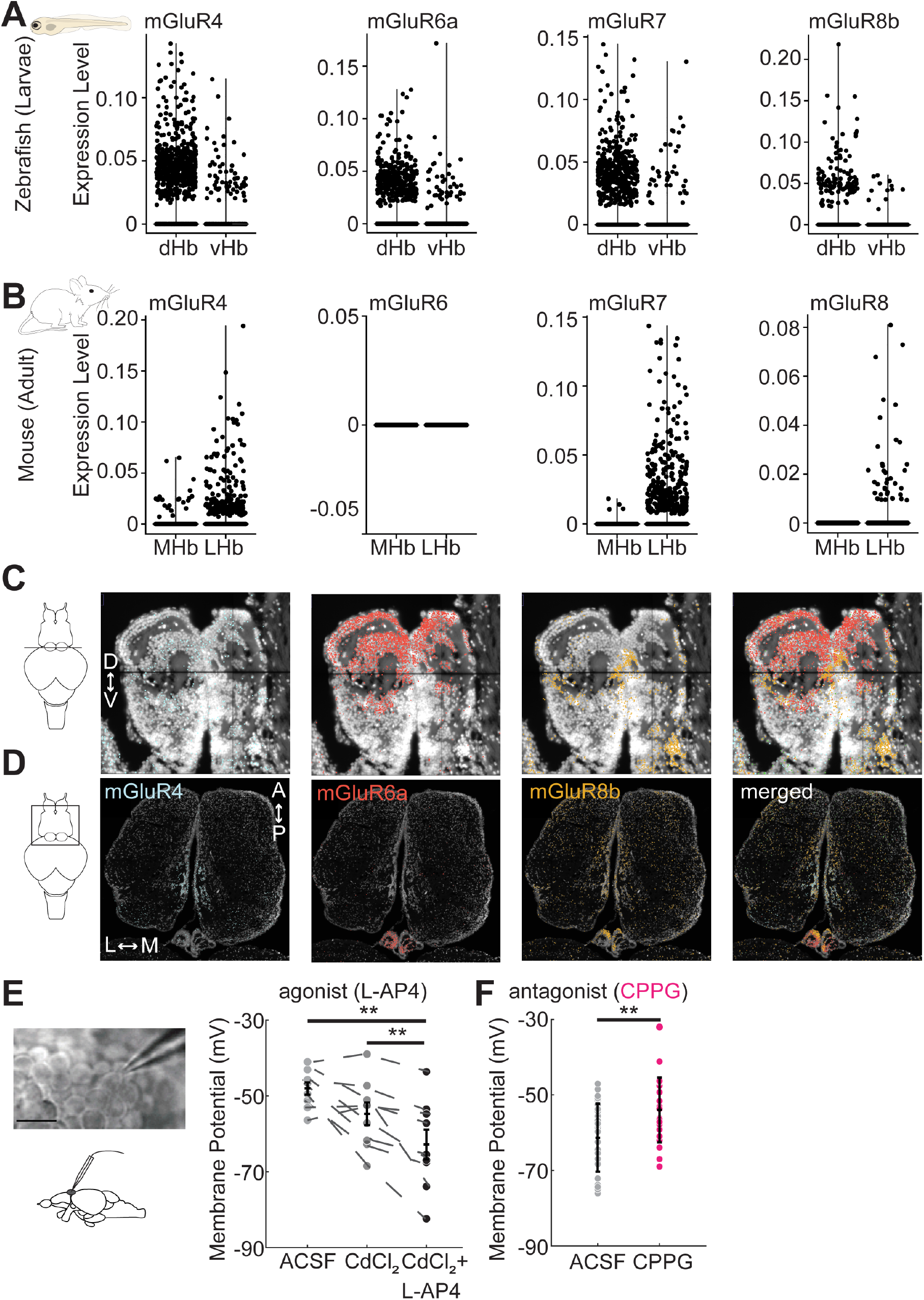
Group III mGluRs are expressed in vertebrate habenula and directly regulate neuronal membrane potential. (A, B) Scatter plots show the mRNA expression of group III mGluRs in zebrafish dorsal (dHb) and ventral (vHb) Hb (A), and mouse medial (MHb) and lateral (LHb) Hb (B). (C, D) High-resolution spatial transcriptomics for mGluR4 (blue), mGluR6a (red), and mGluR8b (orange) in adult zebrafish Hb (C, coronal section), and dorsal forebrain (D, horizontal section), along the dorsal-ventral (DV), anterior-posterior (AP) and medial-lateral (ML) axes. (E) Left: Example image of whole-cell patch clamp recording of a Hb neuron. Scale bar represents 10 μm., Membrane potential measured by whole-cell patch clamp recordings in dHb neurons of juvenile (21dpf) zebrafish brain explants, during control conditions (ACSF), with cadmium chloride (CdCl_2,_ 100 μM) and with CdCl_2_ (100 μM) + L-AP4 (10 μM). Note that group III mGluR agonist L-AP4 decreases the membrane potential of Hb neurons significantly compared to control and CdCl_2,_ conditions. (n=9) p < 0.01, Wilcoxon signed-rank test. Error bars present +/-SEM. (F) Membrane potential of Hb neurons during control condition (ACSF) and with group III mGluR antagonist CPPG (300 μN\M). Note that CPPG significantly increases the membrane potential of Hb neurons compared to control conditions. (ACSF: n = 37, CPPG: n = 22) ** p < 0.01, Wilcoxon rank sum test. Error bars present +/-SEM. See also Suppl. Figure 1.

**Figure 3:**
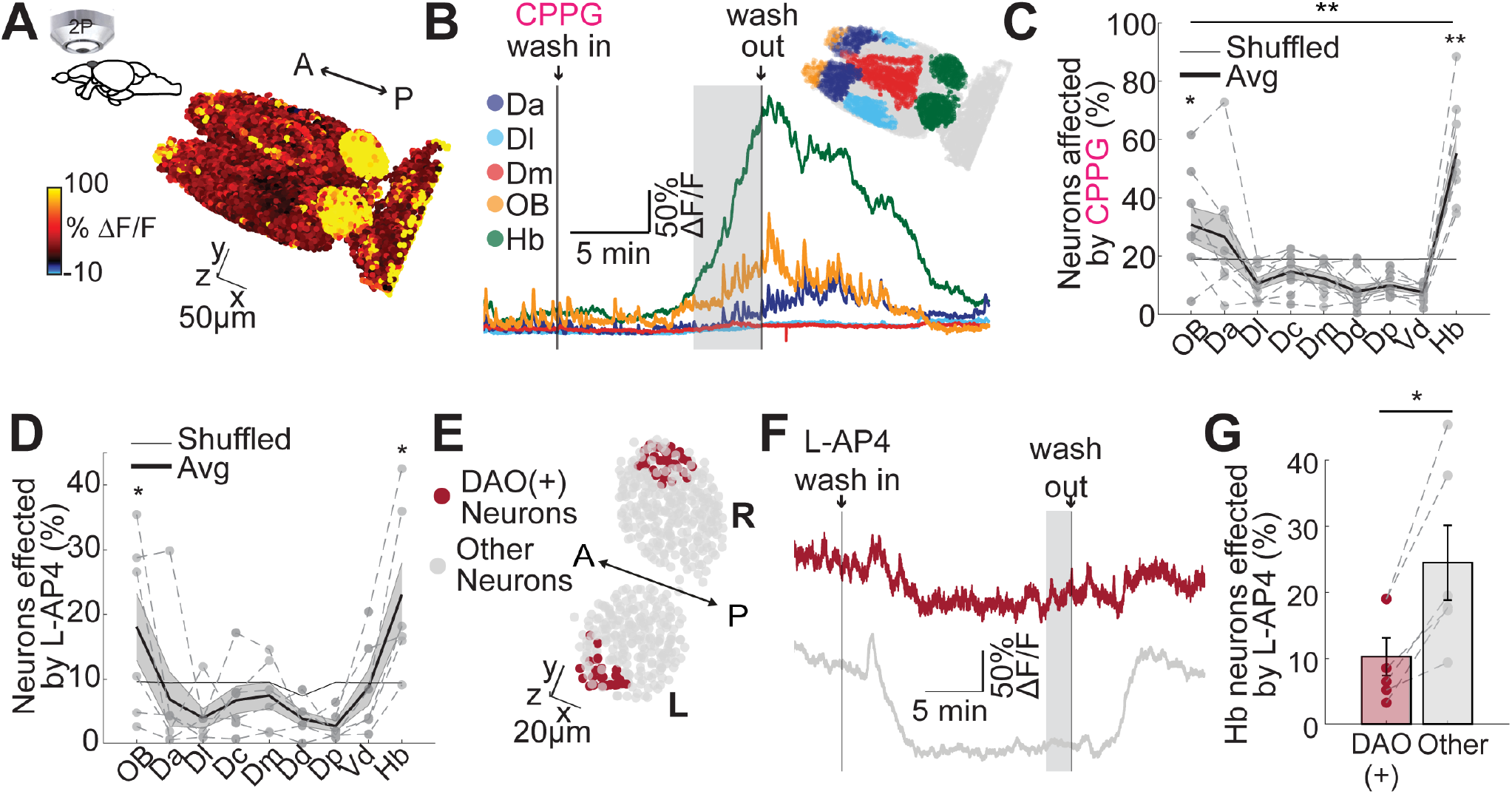
Pharmacological targeting of group III mGluRs can specifically activate and inhibit zebrafish dorsal habenula and the olfactory bulbs. (A) 3D reconstructions of calcium signals (ΔF/F) from a representative juvenile T*g(elavl3:GCaMP6s-nuclear*) zebrafish forebrain in response to bath application of group III mGluR antagonist CPPG (300 μM). Warm colors represent stronger activation. (B) Average time courses of neuronal calcium signals from anatomically identified forebrain regions (Dorso-anterior: blue, Dorso-lateral: cyan, Dorso-medial: red, olfactory bulb: yellow, habenula: green) in response to bath application of CPPG marked by dashed line. (C, D) Percentage of neurons in anatomically identified forebrain region that are affected by CPPG (C) or L-AP4 (0.1 μM) (D). Affected means that during a 5 min drug period (shaded grey in B or F) neuronal calcium signals were 2STD above or below the baseline before drug wash-in. Scatters corresponding to individual fish are connected with dashed lines. Average is indicated as the thick black line; shadow presents SEM. Grey line represents the shuffle distribution. Neurons in the olfactory bulb and the Hb are significantly more affected compared to shuffle distribution (CPPG n = 8, L-AP4 n = 6). Hb neurons show significantly stronger CPPG responses then olfactory bulb. (E) 3D reconstructions of zebrafish Hb expressing both *Tg(dao:GAL-4VP16; UAS-E1b:NTR-mCherry*) and *Tg(eval3:GCaMP6-nuclear*). DAO-positive neurons are indicated in dark red. (F) Average time courses of neuronal calcium signals from example Hb in F for the DAO-positive cells (dark red) and the other Hb neurons (grey). Wash in and out of L-AP4 is indicated by grey lines. The grey area indicated the period in F for affected cell calculation. (G) Significantly smaller fraction of DAO+ vHb neurons are inhibited by the application of group III mGluR agonist L-AP4, when compared to the rest of neurons in dorsal Hb (n = 6). Scatters corresponding to individual fish are connected with dashed lines. Error bars present +/-SEM. L, left; R, right; A, anterior; P, posterior. *p < 0.05, ** p < 0.01, tailed Wilcoxon signed-rank test. Scale bar represents 50 (A) or 20 μm (E).

Next, we asked whether group III mGluRs regulate the membrane potential of dHb neurons in juvenile zebrafish whole-brain explant, by using whole-cell-patch clamp recordings and pharmacological treatments (Figure 2 E&F). To ensure that pharmacological manipulation of group III mGluRs are specific to dHb neurons and not due to effects on neurons presynaptic to the dHb, we first blocked the synaptic transmission by using cadmium chloride (CdCl_2_, 100 μM), a blocker of voltage gated calcium channels. Next, in the absence of synaptic inputs, we applied a group III mGluR agonist L-(+)-2-Amino-4-phosphonobutyric acid (L-AP4, 10 μM). We observed a significant decrease of membrane potential in dHb neurons (Figure 2E), suggesting that group III mGluRs can directly inhibit dHb neurons. In parallel, we also identified that adding (RS)-α-Cyclopropyl-4-phosphonophenylglycine (CPPG, 300 μM), a blocker of group III mGluRs, significantly increased in the membrane potential of dHb neurons (Figure 2F). These results showed that group III mGluRs directly modulate the membrane potential of dHb neurons.

Further, we investigated the impact of group III mGluR agonists and antagonists across the zebrafish forebrain, by using two-photon calcium imaging in *Tg(elavl3:GCaMP6s-nuclear*) 3-weeks-old juvenile zebrafish whole-brain explant (*109, 126-128*). Bath application of group III mGluR antagonist CPPG (300 μM) resulted in an increase of calcium signals mainly in Hb and around the anterior forebrain (Figure 3A). To quantify the distribution of CPPG-activated neurons further, we used anatomical landmarks to delineate forebrain regions (*109, 129, 130*) (Figure 3B). We observed that Hb and the olfactory bulbs are the only forebrain regions with increased calcium signals in a significant fraction of their neurons, with significantly larger fraction in Hb (Figure 3C). In line with this, group III mGluR agonist L-AP4 (0.1 μM) led to significant reduction of calcium signals, in a significant fraction of Hb and the olfactory bulb (Figure 3D).

Our spatial and single cell transcriptomic results revealed that group III mGluR expression is more prominent in dHb as compared to vHb (Figure 2A and C). To test whether the group III mGluRs are less effective in controlling vHb, we performed dual channel two-photon calcium imaging in *Tg(elalv3:GCaMP6-nuclear*) zebrafish where a population of vHb neurons was specifically labeled in red by *Tg(dao:GAL4VP16; UAS-E1b:NTR-mCherry*) (Figure 3E). We identified that the number of DAO-positive vHb neurons responding to the mGluR agonist L-AP4 was significantly smaller than the remaining DAO-negative Hb neurons that are largely in dHb (Figure 3F and G). Altogether, these results revealed that group III mGluRs are present in zebrafish and rodent Hb, and pharmacological manipulation of group III mGluRs can effectively and specifically alter the membrane potential and activity of Hb neurons, with a stronger preference for the dHb.

### Group III mGluRs regulate sensory responses and selectivity of habenular neurons

Our results showed a significant effect of group III mGluRs in Hb excitability and activity. Next, we asked how group III mGluRs regulate multi-sensory representations in zebrafish Hb. To block group III mGluRs *in vivo*, we performed ventricular injection(*108, 131, 132*) of CPPG into *Tg(elavl3:GCaMP6s*) zebrafish larvae (*108, 109, 117-119*). We next investigated the impact of pharmacological group III mGluR perturbation on sensory representations, by comparing Hb responses of animals injected with 5 mM CPPG to control animals injected with artificial cerebrospinal fluid (ACSF) (*128, 133*). We choose two different sensory modalities, relatively neutral red-light flashes (*104, 107, 108*) and aversive mechanical vibrations (*108, 134-137*). To focus on Hb specific effects, we avoided the use of odors due to the expression and impact of group III mGluRs in the OB (Figure 3A-D). We observed that a significantly larger fraction of Hb neurons exhibit excitatory calcium responses to aversive mechanical vibrations in CPPG-injected animals (Figure 4A and B). We also saw a significant reduction of Hb neurons with inhibitory responses to vibrations. This led to an overall increase in average Hb response to vibrations (Figure 4C). Similarly, light responses in Hb also showed a significant increase in CPPG-injected animals (Figure 4D, E and F). The control injections did not alter Hb responses to sensory stimulation (Suppl. Figure 2). Next, we asked whether this significant increase in Hb responses also lead to a reduced selectivity of Hb neurons for these distinct sensory modalities. We observed a significant increase in the number of Hb neurons responding to both sensory modalities (multimodal) in CPPG-injected animals (Figure 4G), with a decrease in unimodal (selective) neurons (Figure 4H). In fact, multi-neuronal sensory representations of vibration and light stimuli were significantly more correlated (more similar) in CPPG-injected animals (Figure 4I). Taken together, blocking group III mGluRs increases sensory-evoked excitation and decreases the selectivity of Hb neurons to vibrations and light stimulation.

**Figure 4:**
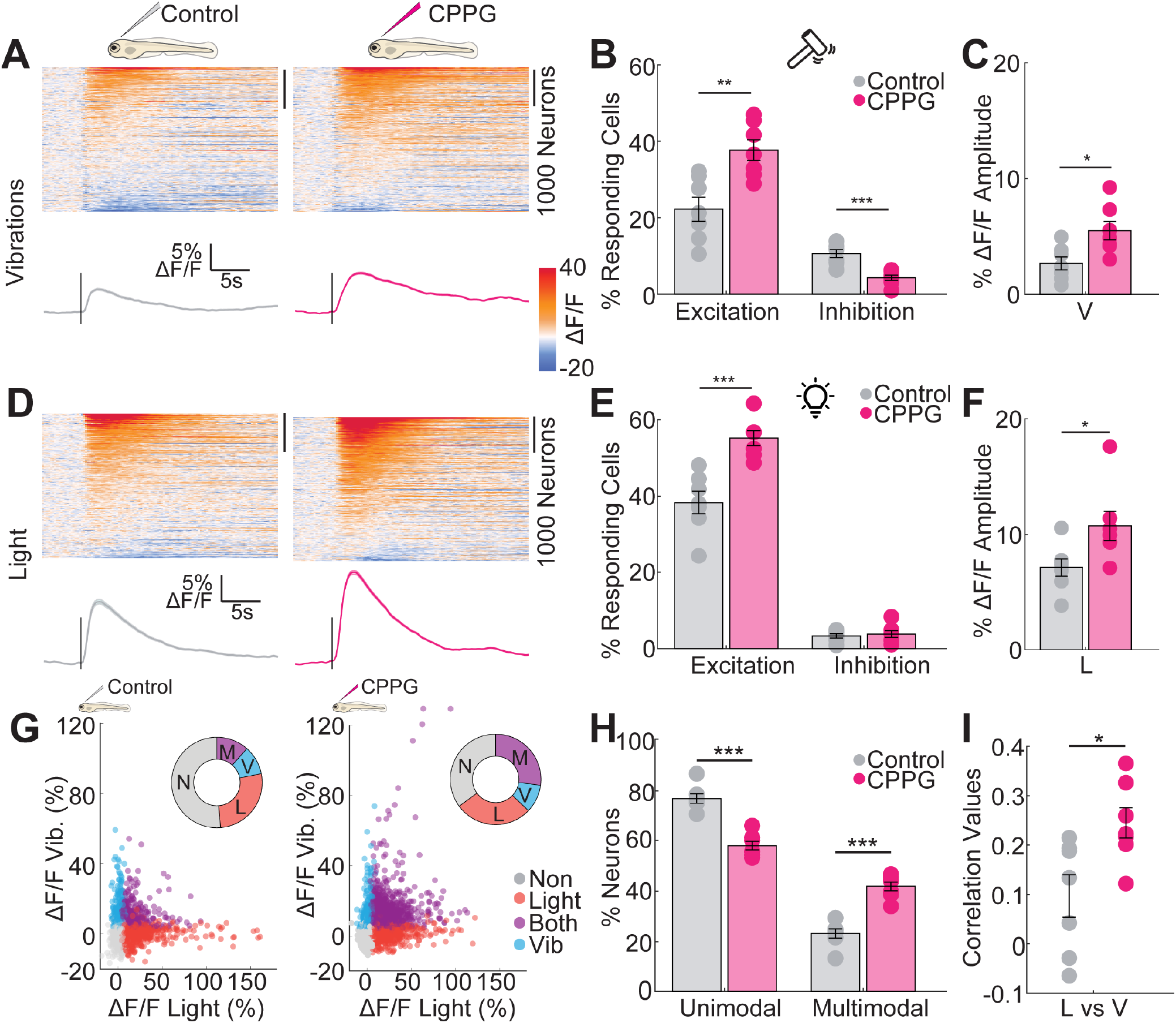
Pharmacological blocking of group III mGluRs amplifies the magnitude and reduces the selectivity of sensory responses of habenular neurons. (A,D) Heatmaps represent the time courses of all Hb calcium signals (ΔF/F) in responses to vibrations (A) or light (D) recorded by two-photon calcium imaging in *Tg(elavl3:GCaMP6s*) larval zebrafish. Left: control-injected (n = 3732 neurons, 7 fish) or right: 5 mM CPPG-injected (n = 4024, 7 fish). Warm colors indicate excitation, cold colors represent inhibition. Average traces of all Hb neurons are below each heatmap. Stimulus onset is indicated by a line. Shadow represents SEM. (B,E) Percentage of excited (2 STD above baseline) or inhibited (1 STD below baseline) Hb neurons for control (grey) or CPPG-injected (pink) fish in response to mechanical vibrations (B) or light (E) stimulation. Note that significantly more cells are excited by both vibrations and light in CPPG-injected fish. Significantly less cells are inhibited by vibrations in CPPG-injected fish. Control n = 7, CPPG n = 7. (C,F) Average ΔF/F Amplitude (%) during the response period of all neurons in Hb per fish. The amplitude after mechanical vibrations or light stimulation is significantly higher in CPPG-injected fish. (G) Responses of individual Hb neurons to mechanical vibration (blue), light (red) or both (magenta) for control and CPPG injected fish. Donut chart represents the ratio of Hb neurons and their response type (2 STD above baseline levels). N: non-responding, V: only vibrations, L: only light, M: both vibrations and light. (H) Percentage of unimodal Hb neurons that respond exclusively to either light (L) or vibrations (V) versus multimodal (M) neurons responding to both light and vibrations. Significantly less neurons in the CPPG-injected fish are selective for one of the two stimulus modalities (unimodal), instead more cells are multimodal. (I) Pearson’s correlation of multi-neuronal response vectors in the Hb for mechanical vibrations and light. Sensory responses in the CPPG-injected fish are significantly more correlated and hence more similar to each other. *p < 0.05, ** p < 0.01, *** p < 0.001, tailed Wilcoxon rank sum test. Error bars present +/-SEM. See also Suppl. Figure 2.

### Inhibitory connections among glutamatergic dHb neurons are mediated by group III mGluRs

Zebrafish Hb neurons were shown to exhibit spatio-temporally structured spontaneous (or resting-state) activity (*106, 108, 109*). Similarly, we observed that positively correlated Hb neurons are closer to each other as compared to negatively correlated neurons in *Tg(elavl3:GCaMP6s*) zebrafish larvae (Figure 5A-B). This can also be observed by a decrease in average pairwise Pearson’s correlations as a function of distance between Hb neurons (Figure 5C, grey). Blocking group III mGluRs by ventricular CPPG injection resulted in an increase in pairwise correlation of spontaneous calcium signals (Figure 5C, pink), suggesting that group III mGluRs regulate the Hb structured activity.

**Figure 5:**
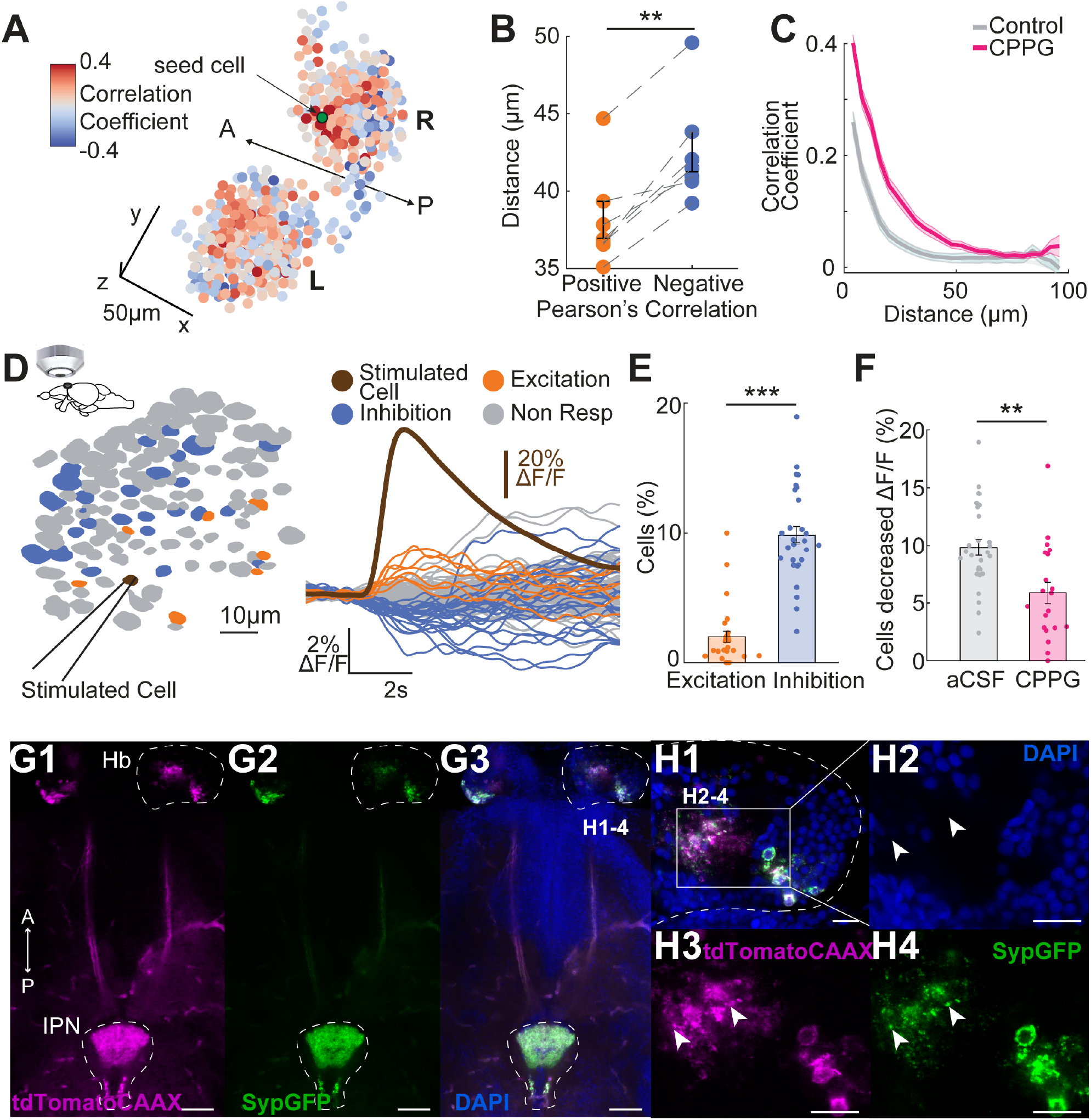
Group III mGluRs play an important role in coordinating spontaneous habenular activity and mediating inhibitory connections between habenular neurons. (A) 3D reconstructions of Pearson’s correlations of spontaneous activity between Hb neurons and an individual seed neuron (green) in *Tg(elavl3:GCaMP6s*) larval zebrafish. Red means high and blue low correlations. Scale bar represents 20 μm) in x, y and z direction. L, left; R, right; A, anterior; P, posterior. (B) Average distance between Hb neuron pairs that are significantly positively or negatively correlated. Note that positively correlated neurons are significantly closer than negatively correlated ones. ** p < 0.01, Wilcoxon signed-rank test (n = 7 fish). (C) Pairwise Pearson’s correlation of Hb neurons during spontaneous activity as a function of distance (μm) between each neuron pair in control (grey) versus 5 mM CPPG-injected fish (pink). Note that CPPG-injected animals exhibit stronger correlations over longer distances. Shadow represents SEM. ANOVA-n displayed significance over distances (** p < 0.01) and over treatment groups (*** p < 0.001). Control: n = 7 fish, CPPG-injected: n = 7 fish. (D) Calcium imaging of *Tg(eval3:GCaMP6-nuclear*) juvenile zebrafish brain explant while simultaneously stimulating a single cell in whole-cell patch clamp configuration. 3D representation of the field of view: brown: stimulated neuron, orange: excited, blue: inhibited, grey: non-responding. Traces of the example neurons are shown on the right (ΔF/F). (E) Percentage of Hb neurons excited or inhibited upon single Hb neuron stimulation. There are significantly more cells being inhibited than excited. *** p < 0.001, Wilcoxon signed-rank test. Error bars present +/-SEM. (F) Percentage of Hb neurons decreasing their fluorescence upon single cell stimulation during control conditions (ACSF) or bath application of CPPG (300 μM). There is significantly less cells being inhibited when 300 μmM CPPG is present in the bath. ** p < 0.01, Wilcoxon rank sum test. Error bars present +/-SEM. (G, H) Confocal microscopy images of *Tg(narp:GAL4VP16*;*UAS: Synaptophysin–GFP-T2A-tdTomato-CAAX*) juvenile zebrafish fixed, tissue-cleared and DAPI stained. Colors represent DAPI (blue), Synaptophysin–GFP (Syp-GFP, green) and tdTomatoCAAX (magenta). (G1) Z-projection confocal image of the entire dorsal Hb-interpeduncular nucleus (IPN) pathway expressing tdTomatoCAAX. White dashed line encircles right Hb zoomed in H1. (G2) SypGFP expression. Note strong expression of Syp-GFP near the axon terminals of Hb neurons at IPN, and the bottom of image panel. (G3) Merged image of tdTomatoCAAX, Syp-GFP, and DAPI staining. Scale bar represents 50 μm). (H1) merged image of zoom into dorsal Hb (encircled with dashed line) from G1-3. White square represents the zoomed area for (H2-4) split into different channels: DAPI (H2, blue), tdTomatoCAAX (H3, magenta), SypGFP (H4, green). Scale bar represents 10 μm). Arrows are pointing at examples of individual SypGFP labelling on the dendritic processes of *narp* labelled dorsal Hb neurons, located in the cell-free (see blue DAPI label) core region of zebrafish Hb. See also Suppl. Figure 3.

The spatial organization of spontaneous activity in zebrafish Hb is likely due to spatially restricted inputs to Hb, as shown for olfactory (*86, 106*), visual (*84, 85, 108*) and hypothalamic (*74*) innervations. Another possible contributor to such functional interactions (correlations and anticorrelations) between Hb neurons are direct connections between them. To test, whether Hb neurons can communicate with each other, we performed patch clamp recordings to stimulate individual dHb neurons, while performing calcium imaging of surrounding Hb neurons in brain explants of *Tg(elavl3:GCaMP6s-nuclear*) juvenile zebrafish (Figure 5D). We observed a strong rise of calcium signals in the micro-stimulated neuron (Figure 5D, brown) followed by an exponential decay. We also observed small but significant excitation (Figure 5D, orange) and inhibition (Figure 5D, blue) in Hb neurons surrounding the stimulated neuron (Figure 5D, E). We observed a larger fraction of Hb neurons which were inhibited (with decreased calcium signals) than excited (with increased calcium signals) (Figure 5E). Blocking group III mGluRs by bath application of 300 μM CPPG, reduced the fraction of Hb neurons that are inhibited upon stimulation of individual Hb neurons (Figure 5F), suggesting that mGluRs may be involved in lateral inhibition.

To investigate the presence of direct connections between Hb neurons, we labelled synaptic terminals of dHb neurons using the *Tg(narp:GAL4VP16*;*UAS:Syp-GFP-T2A-tdTomato-CAAX*) (*63, 120*) zebrafish, where the neural membrane is marked in red and synaptic vesicles in green. The dHb mainly projects to the interpeduncular nucleus (IPN) (*59, 107, 138, 139*) (Figure 5G and Suppl. Figure 3). As expected, we observed that the majority of bright synaptophysin-GFP (SypGFP) expression was at the IPN (Figure 5G2 and Suppl. Figure 3C). However, we also observed substantial SypGFP expression within the core region of dHb (Figure 5H1-4), innervated by Hb neuron dendrites, suggesting that glutamatergic dHb neuron dendrites express presynaptic markers and hence may release neurotransmitters within Hb. Altogether, our findings revealed that glutamatergic dHb neurons can communicate with each other primarily through inhibition, which is mediated, at least in part, through group III mGluRs.

### Multi-sensory stimulus competition in the habenular circuits is partially dependent on group III mGluR

We observed prominent inhibitory interactions within Hb neurons, and an important role of group III mGluRs in such interactions and sensory representations in Hb. We hypothesized that such inhibition across distant Hb neurons might be important for competition between sensory representations, when different sensory cues are presented. To test this hypothesis, we first presented light and vibrational stimuli separately and later simultaneously and measured the sensory responses. Comparing simultaneous or combinatorial delivery of different stimuli to individual light and vibrational responses revealed three different kinds of neurons, super-additive (Figure 6A, orange), depressed (Figure 6A, blue), and neither super-additive nor depressed, which we termed sub-additive. Calculating an interactions index (*140, 141*) for these categories (Figure 6B), revealed that around 60% of Hb neurons showed depressed responses to simultaneous delivery of light and vibrations, when compared to responses to individual stimuli (Figure 6C). Less than 20% of Hb neurons demonstrated super-additive interactions. Hence, most interactions between light and vibrational responses in Hb are depressive (or inhibitory), when both stimuli are delivered simultaneously. Next, we asked to what extent group III mGluR dependent interactions between distant Hb neurons play a role in such depressive interactions between sensory modalities. We observed that ventricular injections of 5 mM CPPG, significantly reduced the fraction of depressive interactions between light and vibrational stimuli, while increasing the amount of sub-additivity. These results showed that competition is the main mode of interaction between sensory representations in Hb and inter-Hb inhibition via group III mGluRs plays a prominent role in such competition.

**Figure 6:**
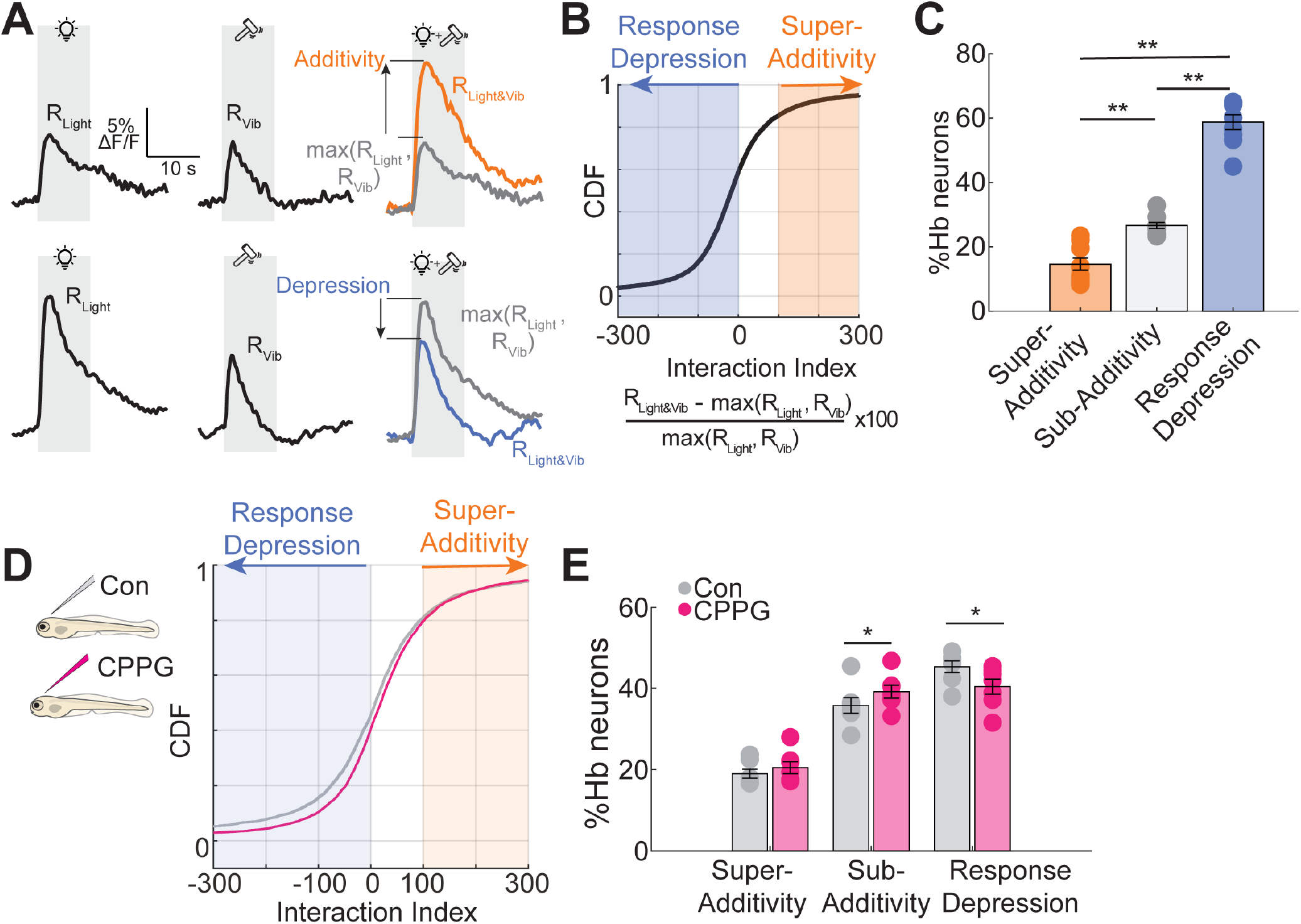
The role of group III mGluRs in multi-sensory stimulus competition in zebrafish habenula. (A) Example traces of Hb neurons responding to light, vibration and light and mechanical vibrations simultaneously. Upper traces show an example of additivity (orange), where combined light+vibration response is larger than the largest individual light and vibration responses (black). Bottom traces show an example of depression (blue) where combined light+vibration response is smaller than the largest of individual light or vibration responses (black). In grey, the maximum response (either light or vibration) is shown. Grey traces near colored traces shows the comparison of combined response to the largest or either light or vibration responses. (B) Cumulative distribution function of the interaction index for each Hb neurons (n = 5242 neurons in n = 9 fish). Values below 0 (blue shadow) are classified as response depression, values above 100 (orange shadow) are classified as super-additivity. Values between 0 and 100 are classified as sub-additivity. Data points above 300 and below -300 are not shown. (C) Percentage of neurons in categories of super-additivity, sub-additivity and response depression after calculation of the interactive index. There are significantly more neurons showing response depression than super-additivity or sub-additivity. (D) Interactive index of Hb neurons from control-injected (grey, n = 3732 neurons in n = 7 fish) and 5 mM CPPG injected-fish (pink, n = 4024 neurons in n = 7 fish). Data points above 300 and below -300 are not shown. (E) Percentage of neurons falling into the categories after calculation of the interaction index. Significantly less neurons show response depression in CPPG-injected fish as well as more cells falling into the sub-additivity category. *p < 0.05, ** p < 0.01, *** p < 0.001, tailed Wilcoxon rank sum test. Error bars present +/-SEM.

### Genetic perturbation of a habenula specific group III mGluR, mGluR6a, alters sensory responses in the dorsal habenula and amplifies defensive behaviors

Previous research has shown that Hb is involved in various defensive behaviors (*57-64, 66, 92, 101-103*). To address the role of group III mGluR mediated inhibition in Hb in sensory-evoked defensive behaviors, we knocked out mGluR6a, which is the main group III mGluR gene expressed in Hb (*142*) (Figure 2D, red), and performed a set of calcium imaging and behavioral experiments. We used 3-weeks-old juvenile zebrafish that can generate cognitively demanding behaviors (*62, 67, 143-147*). We first investigated to what extent mGluR6a knockout recapitulated the pharmacological alterations we observed in the larvae injected with group III antagonist CPPG (Figure 4). Since mGluR6a transcripts are primarily present on dorso-medial Hb (dmHb) (Supp. Figure 4A), we focused our functional analysis on the Hb neurons that are in the most dorso-medial regions (Supp. Figure 4B). Two-photon calcium imaging of juvenile *Tg(elavl3:GCaMP6s*) zebrafish revealed that sensory responses in mGluR6a mutants are significantly elevated (Supp. Figure 4C-H), and less stimulus-specific (Supp. Figure 4I-K). These results are very similar to the changes in sensory representations we observed upon blocking group III mGluRs via ventricular CPPG injections (Figure 4).

Next, we asked whether the behavior of mGluR6a mutant juvenile is altered in well-established defensive behavioral assays. First, we tested mGluR6a mutants in a novel tank test (*148, 149*), where zebrafish express a defensive response by diving at the bottom of the new tank before gradually exploring the tank at various depths (*150-155*). We observed that while control fish initiate such exploratory behavior and swim up, already within first 2 minutes to exposure to new tank, heterozygous and homozygous mGluR6a mutants remained significantly closer to the bottom of the tank (Figure 7A, B). Next, we tested the behavioral response of zebrafish to mechanical vibrations, which are known to generate a reflex-like startle response (*108, 134-137*). We observed that upon mechanical vibrations juvenile zebrafish generate an immediate bottom diving response followed by sustained bottom swimming that outlasts the short (4 seconds) vibrational stimuli for up to 30 seconds (Figure 7C). We observed that mGluR6a mutants remained significantly closer to the bottom of the tank during this sustained behavior. Finally, we used a light-dark transition assay (*156*), where turning off the light generated an initial startle reflex-like response followed by sustained swim with increased speeds in a horizontal arena (Figure 7E). We observed that mGluR6a mutants generate significantly higher sustained swim speeds during light-dark transitions (Figure 7F). In summary, we observed amplified defensive behaviors evoked by environmental changes and sensory stimulation in mGluR6a mutant juvenile zebrafish. Altogether our findings revealed a novel role for group III mGluRs inhibition in Hb in sensory computations, neural connectivity, and animal behavior.

**Figure 7:**
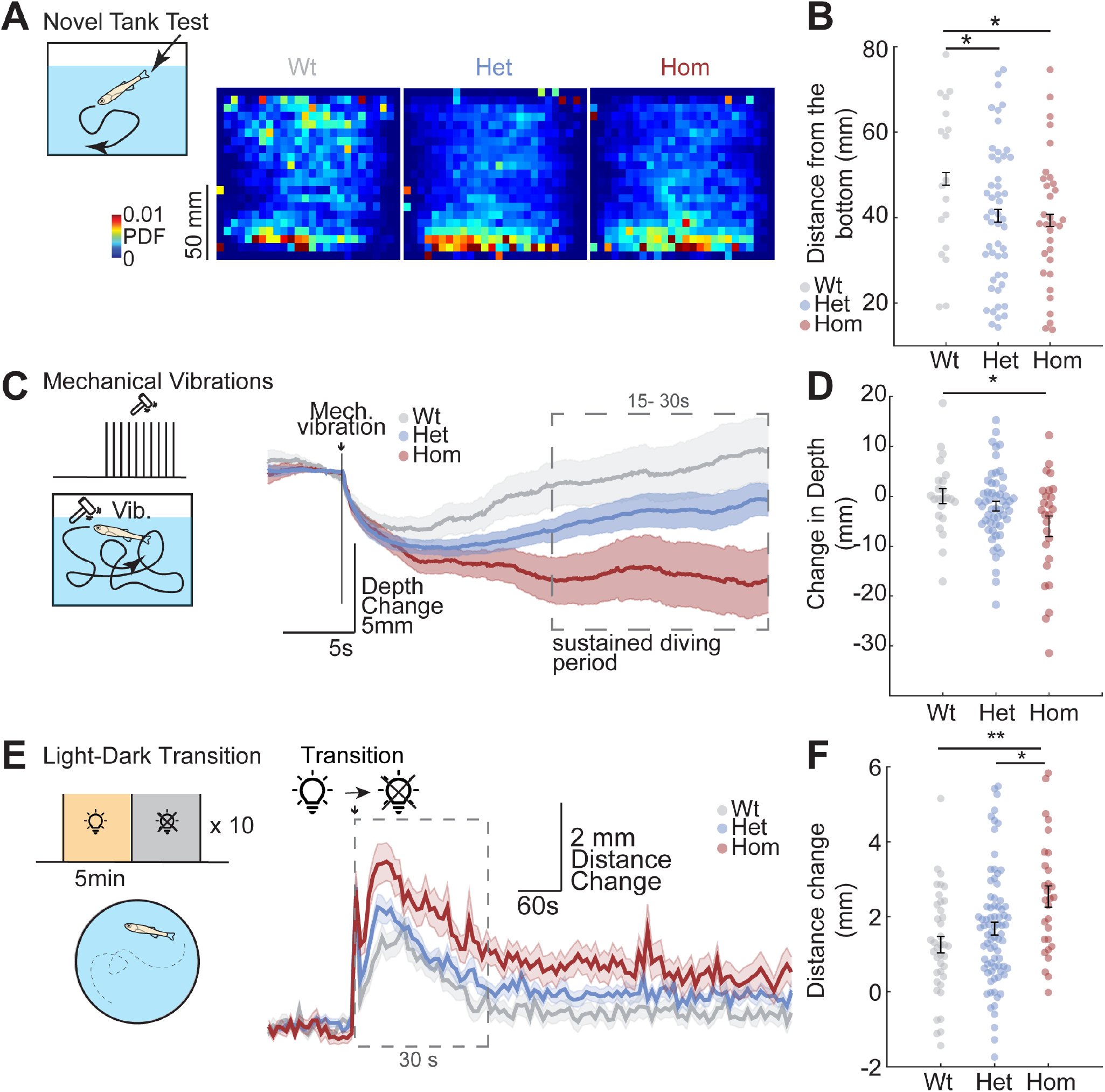
Genetic perturbation of mGluR6a enhances sustained defensive behaviors in juvenile zebrafish. (A) Novel tank diving test of free-swimming mGluR6a knockout, in a vertical tank. Three groups: wildtype (Wt, grey), heterozygous (Het, blue), homozygous (Hom, red). Average probability density of the fish position in the tank, during the first 2 minute after introduction into the novel tank. Warmer colors mean higher probability. (B) Distance from the bottom of the tank (mm) during the first 2 minutes. Note that Het and Hom are significantly closer to the bottom of the tank when compared to Wt fish. Wt n = 18, Het n = 50, Hom n = 32. (C) Mechanical vibrations are delivered to free-swimming juvenile zebrafish, in a vertical tank. Average diving depths evoked by mechanical vibrations, indicated by the grey line. Grey dashed boxes indicate the sustained diving period that was used for statistical comparisons in D. (D) Diving depth during presentation of mechanical vibrations is shown for the three groups. Homozygous mGluR6a mutants have a significant lower change in swimming depths during the sustained diving period marked in C. Wt n = 23, Het n = 54, Hom n = 26. *p < 0.05, Wilcoxon rank sum test. Error bars present +/-SEM. (E) Sustained increase in free swimming speed evoked by Light-Dark transition, in a horizontal tank. Average swimming distances (mm) are plotted for the three groups, the light to drank transition time point indicated the small arrow. Grey dotted boxes indicate the sustained increased swimming rate period evoked by light-dark transition that was used for statistical comparisons in F. (F). Homozygous mGluR6a mutants have a significantly higher sustained increase in free swimming speed (mm) during the dark and light transition. Wt n = 39, Het n = 80, Hom n= 29. *p < 0.05, ** p < 0.01, tailed Wilcoxon rank sum test. Error bars present +/-SEM. See also Suppl. Figure 4.

## DISCUSSION

In this study we investigated the role of group III mGluR-dependent inhibition in brain function and animal behavior. We showed that glutamate-dependent inhibition can mediate several functional features that are usually attributed to GABA, in the Hb, a brain region that largely lacks GABAergic neurons(*108, 110, 112, 116*). By pharmacological and genetic perturbation of group III mGluRs, we showed that Hb neurons exhibit larger and less specific responses to different sensory modalities. We revealed that glutamatergic Hb neurons can directly communicate with each other, largely through inhibition, which is sensitive to group III mGluR blockers. In line with this, we also observed that the default interaction between distinct sensory representations in Hb is mostly based on suppression and depends on group III mGluRs. Genetic perturbation of Hb specific mGluR6a gene, led to amplified defensive responses to sensory stimuli and environmental changes. All these results indicate that group III mGluRs, a rather understudied class of mGluRs in the central nervous system, can mediate features of Hb connectivity and computations that are typically attributed to GABAergic inhibition in other parts of vertebrate forebrain.

The Hb receives inputs from various forebrain regions and sensory systems (*65, 66, 74-80, 84-86, 106-109*). The LHb/vHb receives both glutamatergic (*66, 157-160*) and GABAergic (*66, 78, 157, 161*) inputs. To date, less is known about the GABAergic inputs into MHb/dHb as most reports have studied primarily the glutamatergic system (*162*). Importantly, sensory inputs to the dHb in zebrafish are primarily glutamatergic (*84-86, 120*) and MHb/dHb neurons are glutamatergic as well (*108, 110*). Despite this, sensory-evoked inhibition is present in dHb (*106-108*). Hence, the GABA-poor dHb appears to have evolved an inhibitory system that does not require GABA. We now showed that inhibition in dHb at least in part relies on glutamate. Glutamatergic excitation and inhibition of Hb neurons might be acting on different time scales. Upon sensory stimulation (*106-108*) glutamate release in dHb can generate a fast excitation via NMDA/AMPA receptors on Hb neurons (*109, 112, 116*). Whereas mGluRs dependent inhibition is slow, since it relies on multiple cellular cascades (*49-51*). Moreover, group III mGluR are thought to be present both in the synapse and extra synaptically (*49, 163*). Such extra-synaptic localization might also contribute to slower dynamics of mGluR dependent inhibition. Altogether these results propose that mGluR dependent inhibition has slower temporal kinetics, when compared to fast glutamatergic excitation. Hence, glutamate release in Hb can generate a fast excitation followed by slowly spreading inhibition, and therefore mGluR-dependent inhibition in Hb can be important for slow adaptive responses and behaviors, as we have observed in juvenile zebrafish (Figure 7).

We showed that suppression is the default integration mode of Hb neurons for distinct sensory inputs, which resembles lateral inhibition in sensory systems (*9, 11, 164-168*). Such cross-modal inhibition is in line with the proposed function of Hb to act as a switchboard (*57, 76, 106, 108, 143*). In fact, our results suggest that suppression across different modalities as well as Hb ensembles might be the primary substrate for such a switchboard function of Hb. This cross-modal inhibition in Hb is at least partly mediated by group III mGluRs receptors in GABA-poor dHb/ MHb, as well as by multi-synaptic inhibition through Hb-interpeduncular nucleus (IPN) connections onto presynaptic terminals of Hb neurons (*115*). Both dHb/MHb and vHb/LHb are instrumental for adaptive behaviors (*57-64, 66, 68, 92, 101-103*) and mood disorders (*71-73*). We observed only minimal expression of group III mGluRs in zebrafish vHb, which was shown to receive GABAergic terminals that likely mediate most observed inhibition in vHb (*169*). However, prominent expression of group III mGluRs in rodent LHb suggest that such inhibition can still play unknown roles in mammalian LHb. Coincidentally, few studies showed noticeable effects on defensive and anxiety-like behaviors in models of mood disorders, upon perturbation of group III mGluRs, mGluR4 and mGluR8 in various mammalian forebrain regions (*52, 170-177*). Given relatively specific responses of zebrafish Hb to group III mGluR pharmacology, we proposed that group III mGluRs can be a potential drug target for interfering with Hb function and mood disorders.

In our study, we observed several neural circuit features such as coordinating ensembles (*113, 178, 179*), cross modality suppression (*24, 140, 180*), sensory selectivity and gain (*10, 12, 14, 18, 20, 23*) that are attributed to GABAergic circuits across the vertebrate forebrain, are at least in part mediated by group III mGluRs in dHb. Yet, group III mGluRs are vastly expressed throughout mammalian brain (*50, 52, 53, 181, 182*). While few studies investigated the function of group III mGluRs in retina (*50, 183, 184*), we know very little about the role of mGluR-dependent inhibition in brain function and animal behavior. An interplay between group III mGluRs and GABAergic interneurons has been seen in a recent study (*185*), which suggests glutamatergic and GABAergic inhibition may complement each other. Recently available molecular atlas of mammalian brain (*54, 186, 187*) suggests broad expression of group III mGluRs such as mGluR7 and mGluR8 in various cortical areas, from entorhinal to sensory/motor and associative cortices. Interestingly, group III mGluRs were suggested as drug targets for various human disorders like neurodegenerative diseases (*50, 188*), anxiety (*170*) and absence epilepsy (*189*). Our results in Hb and these previous studies across the brain highlight the potential of studying group III mGluRs function from the perspective of brain physiology and pathophysiology. Future studies will reveal what other functions glutamate driven inhibition may play in vertebrate brain.

## ACKNOWLEDGEMENTS

We thank M. Ahrens (HHMI, Janelia Farm, USA), K. Kawakami (NIG, Japan), and H. Okamoto (RIKEN-CBS, Japan) for transgenic lines; Y. Yoshihara RIKEN-CBS, Japan for sharing the *UAS:Syp–GFP-T2A-tdTomato-CAAX* plasmid, S. Eggen, M. Andresen, V. Nguyen, and our fish facility support team for technical assistance; the Yaksi lab members for stimulating discussions. This work was funded by ERC starting grant 335561 (E.Y.), NFR FRIPRO research grants 239973, 314212 (E.Y.) and 314189 (N.J-Y), an RSO grant from Norwegian University of Science and Technology (E.Y, A.M.O.); and Boehringer Ingelheim Fonds (S.K.J.). Koc University Neuroscience Master’s Program fellowship (YIC). Work in the E.Y. laboratory is funded by the Kavli Institute for Systems Neuroscience at Norwegian University of Science and Technology.

## AUTHORS CONTRIBUTIONS

Conceptualization, A.M.O. and E.Y.; Methodology: A.M.O., N.F., B.S., I.J., A.E., F.H., R.B., S.K.J., N.J-Y. Analysis A.M.O., N.F., Y.I.C.C., A.K.M., and E.Y.; Investigation, all authors; Writing A.M.O., and E.Y.; Review & Editing, all authors; Funding Acquisition and Supervision, E.Y.

## SUPPLEMENTARY INFORMATION

Document S1: Figures S1-S4

## METHODS

### Zebrafish maintenance and strains

The animal facility and fish maintenance were approved by the NFSA (Norwegion Food Safety Authority). Zebrafish, *Danio rerio*, were kept in 3.5 L tanks in a Techniplast ZebTec Multilinking System with constant conditions maintained (28.5 C, pH 7.2, 700 uSiemens, 14:10 h light/dark cycle). The fish were fed with dry food (SDS100 up to 14 dpf and SDS 400 for adult animals, Tecnilab BMI, the Netherlands) twice a day and *Artemia nauplii* (Grade 0, Platinum Label, Argent Laboratories, Redmond, USA) once a day. Larvae from fertilization to 3 dpf were kept in a Petri dish with egg water (1.2 g marine salt in 20 L RO water, 1:10000 0.01% methylene blue) and in artificial fish water (AFW: 1.2 g marine salt in 20 L RO water) between 3 and 5 dpf. Larval (5dpf) and juveniles (3 to 4-week-old) zebrafish were used for the experiments and analyzed irrespective of gender. The following fish lines were used for experiments: *Tg(elavl3:GCaMP6s*)(*119*), T*g(elavl3:GCaMP6s-nuclear*) (*119*), *Tg(dao:GAL4VP16; UAS-E1b:NTR-mCherry*)(*58*),*Tg(narp:GAL4VP16*;*UAS:Syp–GFP-T2A-tdTomato-CAAX)(63, 120*) mGluR6a mutant was created using a CRISPR/Cas9 mediated knockout.

### Ethical guidelines statement

All experimental procedures that were performed on zebrafish larvae and juveniles were in accordance with the Directive 2010/63/EU of the European Parliament and the Council of the European Union. They were approved by the Norwegian Food Safety Authorities.

### Two-photon calcium imaging

For calcium imaging two photon microscopes (Scientifica) in a combination with a 16x water immersion objective (Nikon, NA 0.8, LWD 3.0) and a 20x water immersion objective (Zeiss 7 MP, W Plan-Apochromat NA 1.0) were used. A mode-locked Ti:Saphire laser (MaiTai Spectra Physics) tuned to 920 nm was used for excitation. In addition, recordings were either performed single plane or volumetric (8 planes using a Piezo (Physik Instrumente (PI))). Volumetric recordings had an acquisition rate of 2.3-3.4 Hz per plane and an average images size of 1536 × 600 pixel.

Fish for *in vivo* imaging were embedded in low melting point agarose (2-2.5 % LMP, Fisher Scientific) inside a recording chamber (Fluorodish, World Precision Instruments) for head-restraining. After 20 min the LMP agarose solidified and the section covering the mouth and nose was carefully removed for juvenile zebrafish but left intact for larval ones. The embedded fish was then placed under the microscope and in case of the juvenile zebrafish constantly perfused with bubbled AFW. Before any sensory stimulations ongoing activity was always recorded for 10 min. A red LED (LZ1-00R105, LedEngin; 625-nm wavelength) was used for light stimulations and placed it in the front of the recording chamber. The duration of the light stimulus was a flash of 2 ms with an intensity of 0.318 mW. Mechanical vibrations were delivered via solenoid tapper (SparkFunElectronics, ROB-10391), via 50-ms application of 6 V. For co-stimulations of the two sensory stimuli, the onsets were synchronized. Light, vibration or co-stimulation of light and vibration were repeated eight times with a 60 s interstimulus interval, and 4 min were between the different sets of stimulus conditions. For odor stimulation we prepared an amino acid mixture (Alanine, Phenylalanine, Methionine, Histidine, Cysteine, Arginine, Glutamic acid at 10^−4^ M), and ammonium chloride (10^−4^ M). All odorants were purchased from Sigma Aldrich. The odor was delivered via stimulation tube that was placed in front of the nose of the fish. The stimulation lasted 30 s and was performed using a HPLC injection valve (Valco Instruments) controlled with Arduino Due. To determine the precise onset of odor delivery a trial with fluorescein (10^−4^ M AFW) was performed before each experiment.

For the whole brain explant imaging, the dissection of the juvenile zebrafish explant was described previously(*109*). In short, animals were anesthetized in ice-cold AFW and euthanized by decapitation in oxygenated (95% O_2_/5% CO_2_) ACSF. The ACSF was composed of the following chemicals diluted in reverse osmosis-purified water: 131 mM NaCl, 2 mM KCl, 1.23 mM KH2PO4, 2 mM MgSO47H2O, 10 mM glucose,

2.5 mM CaCl2, and 20 mM NaHCO3 as previously explained (*128*). Both, the dissection, and experiment with the brain explant was carried out in bubbled ACSF. After removing the jaw and eyes, the muscle tissue, gills, and fat were cleaned of to ensure proper oxygen diffusion of the explant. Then, bones, skin tissue and dura mater covering the forebrain and/or Hb were removed from the dorsal side. The brain explant was then mounted using tungsten pins to a small petri dish coated with Sylgard (World Precision Instruments) and perfused in constant flow bubbled ACSF. The brain explant is consistently perfused with artificial cerebrospinal fluid (ACSF) that is bubbled with carbogen (95% O2 and 5% CO2) throughout the entire duration of the experiment.

### Electrophysiological recordings

Patch clamp recordings were conducted in a juvenile brain explant. For intracellular recordings of Hb neurons, borosilicate glass capillaries of 9-15 MOhms were filled with intracellular solution which contained (in mM):

130 K-gluconate, 10 Na-gluconate, 10 HEPES, 10 Na^2+^-Phospho-Creatine, 4 NaCl, 4 ATP-Mg and 0.3 Na^3+^-GTP (*128*). Electrical signals were recorded by MultiClamp 700B amplifier at sampling rate of 10 KHz. All recordings and data analyses were performed using custom codes written in MATLAB. To perturbed group III mGluRs the agonist, L-AP4 (Tocris) with a concentration of 10 μM and the antagonist, CPPG (Tocris) with a concentration of 300μM was used. To block synaptic transmission cadmium chloride (100μM) was used. Single cell microstimulation were performed in juvenile brain explains (see “Explant Preparation”). With a borosilicate glass capillaries neurons were patched (9–15 MOhms) and then stimulated using 0.04-0.5 mA current injections for 500 ms. A cell was stimulated with up to 60 pulses in total, stimulating 6 times per sweep with 10 sweeps except for one cell that has 7 sweeps (42 pulses in total). To block group III mGluRs, CPPG with a concentration of 300μM was used. The calcium activity of Hb was recorded using the 10x objective in the green channel at a frame rate of 10 frames per second.

### Confocal and anatomical imaging

For confocal imaging 21 dpf, *Tg(narp:GAL4VP16*;*UAS:Syp–GFP-T2A-tdTomato-CAAX*) fish were used. Brains were dissected similar to the explant preparation and then fixed and cleared. In addition, samples were stained with DAPI (4′,6-diamidino-2-phenylindole, 1:1000 for 2h). Anatomical Z-scans were acquired using a Zeiss Examiner Z1 confocal microscope with a ×20 plan NA 0.8 objective or x40 plan NA 1.4 oil objective. For the example confocal image of the *Tg(narp:GAL4VP16*;*UAS:Syp–GFP-T2A-tdTomato-CAAX*) brains were cleared using a clearing kit (Binaree) and stained with 4’,6-diamidino-2-phenylindole (DAPI). After fixation, the zebrafish brains were cleared using the Binaree Tissue ClearingTM Kit (#HRTC-012, Binaree, Republic of Korea) using the recommended protocol. On the 3^rd^ day of the clearing process, the brains were stained with 1:3000 dilution DAPI for 12h at 4ºC before being washed with distilled water and then transferred to the mounting solution provided by the kit (#HRMO-006, Binaree, Republic of Korea). The brains were then kept at room temperature (RT) until imaging.

For the RNA detection of mGluR6a we used hybridization chain reaction (HCR) on juvenile zebrafish brains previously described (*190*). HCR is combined with a shorten version of the Binaree’s clearing protocol. Juvenile zebrafish were euthanized in cold AFW. After euthanasia, juvenile zebrafish were transferred into a 1.5 mL tube containing 200 μL of 4% PFA in dPBS and incubated at 4 °C overnight. Brains were dissected out after washing the samples with 1X dPBS three times at RT for 5 minutes. Then, the samples were dehydrated and permeabilized with 100% methanol (MeOH) washes for 10 minutes 4 times and 50 minutes 1 time before getting stored at -20°C overnight. The next day, the samples were rehydrated with a series of graded 1 mL MeOH/dPBS washes for 5 minutes each at RT: (a) 75% MeOH in dPBS, (b) 50% MeOH in dPBS, (c) 25% MeOH in dPBS, and (d) dPBS. For the clearing stage, the Starting solution from Binaree is used to incubate the samples at 4°C overnight. Next day, 500 μL of Tissue Clearing Solution B is added and incubated in a water bath at 37°C for 24 hours. Then, the samples were washed with reverse osmosis water while shaking at 30 rpm at 4°C for 30 minutes, repeated four times. 500 μL probe hybridization buffer is added, and samples are incubated for 30 minutes at 37°C. After removing the probe hybridization buffer, a probe solution (2 pmol of each probe set, prepared by mixing 2 μL of 1 μM stock in 500 μL of probe hybridization buffer at 37°C) is added. The samples were incubated overnight at 37°C and washed four times for 15 minutes each with 500 μL of probe wash buffer at 37°C. Lastly, the samples are washed twice for 5 minutes each with 500 μL of 5x SSCT at RT. 500 μL of amplification buffer is added and incubated for 30 minutes at RT for the amplification stage. Meanwhile, 30 pmol of hairpin h1 and 30 pmol of hairpin h2 are separately prepared by snap cooling (The tubes are heated to 95°C for 90 seconds and then cooled to RT in a dark drawer for 30 minutes) 10 μL of 3 μM stock. The snap-cooled h1 and h2 hairpins are added to 500 μL of amplification buffer at RT, and the samples are transferred into this mixture and incubated overnight in the dark at RT. Samples are washed with 500 μL of 5x SSCT at RT as follows: (a) 2 × 5 minutes, (b) 2 × 30 minutes, (c) 1 × 5 minutes, and with dPBS for 3 × 5 minutes. The samples are then transferred to 500 μL of the Mounting Solution from Binaree and incubated overnight (12–16 hours) in the dark at RT. LSM880 upright Zeiss confocal microscope with 20X (Numerical Aperture: 0.8, Working Distance: 0.55 mm) objectives is used for imaging the samples.

### Generation of *Tg(5xUAS:Syp-GFP-T2A-TdTomato-CAAX*)^*nw19Tg*^ line

To generate the transgenic line, the *UAS:Syp–GFP-T2A-TdTomato-CAAX* plasmid DNA was used (*120*). 2 nL of a mixture of the plasmid DNA (60 pg) and tol2 mRNA (10 pg) was microinjected into one-cell stage embryos, as previously described in Jeong et al. (*132*). The injected embryos were raised to adulthood (F0). Germline-transmitted founder zebrafish were identified by breeding with several Gal4 transgenic lines. Stable F1 larvae showing the Gal4 driven GFP and membrane-tethered TdTomato signals were raised to adult zebrafish.

### Ventricular Injections

Larval zebrafish (5 dpf) were mounted in 2% LMP agarose. After solidification of the agarose, the fish were injected with either 1 nL CPPG (5mM) or vehicle (NaOH, 5mM) using a glass pipette into the forebrain ventricle. Ventricular injections have been previously described (*108, 131, 132*). After a waiting period of 10 min, the 2P imaging experiment was conducted as explained in section “Two photon calcium imaging”.

### Generation of the mGluR6a mutant

Crispr/Cas9 mutagenesis was performed as previously described in Schlegel et al. (*191*). CRISPR target sites of the mglur6a gene selected using CHOPCHOP (chopchop.rc.fas.harvard.edu). Targets sites were: CGAGGAGGTCCAATCTAACC (T7) and GACCAGGAGGACGTGGCTGA (Sp6).

Forward and reversed oligonucleotides (pT7-gRNA_fwd: CAGCTATGACCATGATTACG; pT7-gRNA_ rev: AAAAGCACCGACTCGGTG; pSp6-gRNA_fwd: ATTTAGGTGACACTATA; pSp6-gRNA_rev: ATTTAGGTGACACTATA) were cloned into pT7- or pSP6-sgRNA as described in Jao et al. (*192*). sgRNAs were transcribed using MEGAscript Sp6 kit (Ambion, Austin, TX, USA) and MEGAshortscript T7 kit and purified using using the MEGAclear kit (Ambion, Austin, TX, USA). The resulting mutant has a 103bp deletion (alignment by request) with no detectable protein labeling (data not shown).

### Behavioral assay

The novel tank test was conducted in a custom behavioral set up that allowed to simultaneously record 6 fish in 6 separate arenas (11.5 cm x 11.5 cm x 1.5 cm) from the side. In short, fish were introduced to the arena in the first 5 seconds of the experiment where the bottom row of fish was introduced first, following the top three fish. The fish were then tracked for their x and y position and the data later analyzed with custom MATLAB scripts. The total length of the experiment was 30 min. The time point used for the analysis was the first two minutes. The mechanical vibration assays were conducted in an Zantiks LT set up with 6 arenas (10 cm x 11.5 cm x 3 cm) allowing to perform the experiment with six fish simultaneously while recording their position from the side. The fish were individually tracked (x and y position) and the data later analyzed in custom MATLAB scripts. For the mechanical vibration experiment, the fish were habituated to the arena for 20 min. Then after a 4 min baseline, 15 mechanical vibrations were delivered to the 6 arenas simultaneously with an interstimulus interval of 1 min, followed but a 4 min baseline period. The Light/Dark Transition experiment was conducted in an Zantiks MWP set up using 6-well plates (VWR) that allow tracking of the animal movement from above. In short, fish were introduced to the well plates and placed in the set up. After a habituation period of 15 min in light, the behavioral protocol started with the dark condition (5 min) and then back to the light condition (5min). This was repeated 10 times. The fish were individually tracked (x and y position) and the data later analyzed in custom MATLAB scripts.

Experiments conducted with juvenile mGluR6a mutants were done blindly by using a heterozygous incross. After the experiment, the fish were genotyped using real-time quantitative polymerase chain reaction melt-curve analysis (*193*). Using the forward primer GCTTGAGCATAAAACTCTAATTC and the reverse primer CAGAGGATGCACATTATATTTC.

### Transcriptomics analysis

The scRNAseq datasets from zebrafish larvae (*112*) and mouse (*116*) were analyzed using R and the Seurat R Package v3 (*194*). Non-neuronal cells in larva were removed based on the expression of *snap25a*, for mouse they were removed based on the expression of *stmn2, thy1* and *snap25*. To identify dorsal and ventral Hb in the zebrafish, *kiss1* and *aoc1* were used, respectively. For the mouse MHb was identified with *tac2* and LHb was identified with *pcdh10* marker genes. Molecular cartography a multiplex method from Resolve Bioscience was performed in adult zebrafish whole brain coronal and horizontal slices with probes for mGluR4, mGluR6a, mGluR6b and mGluR8b. A more detailed protocol is described in D’Gama et al. (*124*).

### Ages of fish

Juvenile zebrafish have been shown to perform cognitively demanding behaviors (*62, 67, 143-147*) which is why we used this age group for most of our experiments. 3-4 weeks old juvenile zebrafish were used for preparing the whole-brain explant in electrophysiology and pharmacology experiments, as well as *in vivo* calcium imaging of mGluR6a mutants and all behavioral experiments. For the *in vivo* ventricular injections, larval zebrafish (5 dpf) were used due to national ethical limitations in performing such *in vivo* ventricular injections in juvenile zebrafish.

### Data Analysis

Whole brain explant two-photon microscopy images were aligned using methods described in previous work (*108, 109, 117, 118*). The *in vivo* recordings were aligned using suite2p (*195*). After alignment all the recording were treated the same. The recordings were visually inspected for any remaining motion z-directional drift and discarded if any motion artifacts remained. Using a template matching algorithm (*106, 109, 118*), regions of interests (ROIs) corresponding to neurons were detected and visually confirmed. Pixel belonging to each ROI were averaged over time to calculate the time course of each neuron. Then the fractional change in fluorescence relative to the baseline (ΔF/F) was calculated. For sensory stimulation trials, the signal was normalized to a 5s pre-stimulation period. For the spontaneous period *in vivo* and the whole brain explant experiments an average over moving window of 6 min was used. For the *in vivo* recordings the calcium signals were smoothed using filtering previously described (*126*).

For the calcium imaging while stimulating a single cell, the 10 (or 7) sweeps were combined into a continuous recording and then motion corrected using NoRMCorre (*196*). Recordings with significant motion artifacts observed after manual inspection were discarded. In the remaining recordings, a ROI was drawn around the Hb neuron that was patched. The rest of the cell detection as well as the signal extracted using CNMF-E (*197*). In addition, the cell detection accuracy was manually inspected and detected ROIS that did not relate to an actual cell were rejected. The patched neuron was manually isolated from the rest of the neurons. The extracted calcium traces were smoothed using a gaussian convolution and normalized using the mean and baseline from the pre-stimulation period.

All the following analysis was performed in MATLAB.

Responding cells (light or mechanical vibrations) were calculated by comparing the response average over a 10s-time window after stimulus onset to the baseline activity. Excitatory responses exceed two times the standard deviation (STD) plus the mean of the baseline duration. Inhibitory responses are below the STD minus the mean of the baseline period. For the single cell stimulation experiments, a not tailed Wilcoxon signed-rank test was performed using median values from the pre-stimulation period and the first two seconds post-stimulation. If a neuron was responsive (p < 0.05), it was considered inhibited if the median of the post-stimulation period was below the median of the pre-stimulation period and excited if the post-stimulation median was above the pre-stimulation median. Recordings in which the stimulated cell did not show statistically significance (excited) were rejected.

Affected cells to the antagonist (CPPG) and agonist (L-AP4) were calculated in a similar way, with CPPG-affected neurons are exceeding two times the STD plus the mean of the baseline period (5 min before wash in) during the drug period (minute 13 to 18 of the experiment) and L-AP4-affected neurons responses are smaller than the mean minus the STD of the baseline period (2 min before wash in) during the drug period (minute 26 to 28 of the experiment).

Delineation of the forebrain regions was done manually according to anatomical landmarks as described previously (*109, 129, 130*).

The response vectors to light or mechanical vibration were vectors of the average response of each neuron during the response period. To find their similarities the two vectors were correlated with each other using Pearson’s correlations.

To understand the response observed in Hb neurons to the combined presentation of mechanical vibration and light, we calculated the interaction index which is based on previous work (*140, 141*) and also knowns as the “interactive index” in the original publication. It is calculated as followed:

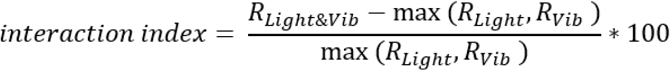

The responses (R) to light, mechanical vibration or the co-stimulation (light and vibration) are average over the response period (10s after stimulus onset). An interaction index of above 100 represents an “super-additive” response where the response to the co-stimulation is at least twice as big as the maximal response to the individual stimuli. On the other hand, a interaction index of below zero represents a response to the co-stimulation that is smaller than the max response of the individual stimulation called “response depression”. An interaction index between 0 and 100 is named “sub-additivity”.

To find the most dorsomedial Hb neurons for mGluR6a mutant imaging experiments, the y-position of the neurons was normalized to the middle point (mean). The z-position was normalized to the highest point. Neurons were then ranked according to the absolute sum of the normalized y and z position. Small sums represent neurons that are dorsal and close to the midline. The 40% percent neurons with the smallest sum were classified as the most dorsomedial neurons.

For the behavioral experiments the x and y positions of the animals were tracked. The heatmaps represent the average density of zebrafish positions. To calculate the distance from the bottom, the y-position of the animals were normalized to the dimensions of the tank. The change in depth was calculated by fraction of the y-position two seconds before the mechanical stimulation and the y-position after the stimulation. The distance change is the fraction of the swimming distance of the fish in x-y direction in 1 s bins five seconds before the transition and the rest of the duration.

### Quantification and statistical analysis

MATLAB was used for the statistical analysis. P-values are represented in the figure legends as * p<0.05, **p<0.01, ***p<0.001. All analysis was performed in MATLAB, R and Fiji.

