## Supplementary Figures 1-4 for "Inhibition mediated by group III mGluRs regulates habenula activity and defensive behaviors"

### Supplementary Material

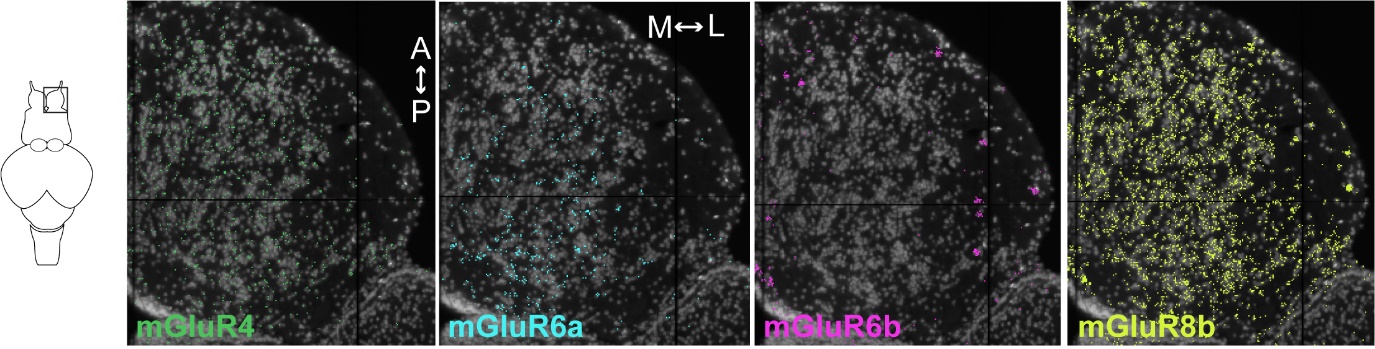

Supplementary Figure 1: Group III mGluR expression in the olfactory bulb of adult zebrafish. High-resolution molecular cartography of the left adult zebrafish olfactory bulb illustrating the expressions of mGluR4 (green), mGluR6a (cyan), mGluR6b (pink) and mGluR8b (yellow). Horizontal section displaying gene expressions along the anterior-posterior (AP) and medial-lateral (ML) axes. AP, anterior-posterior axis; ML, medial-lateral axis; VD, ventral-dorsal axis.

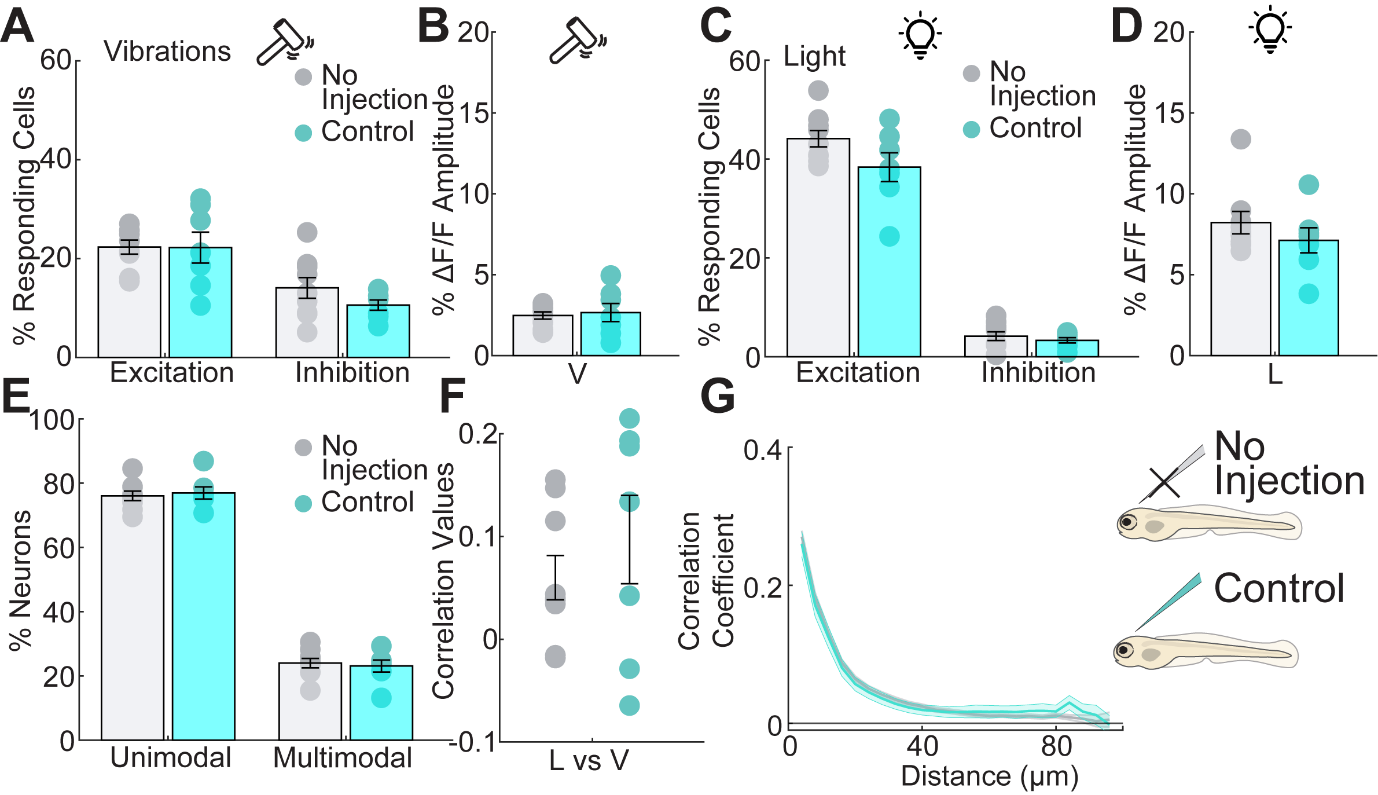

Supplementary Figure 2: Comparing sensory responses and selectivity of non-injected and control-injected fish. Larval *Tg(elavl3:GCaMP6s)* zebrafish recorded by two-photon calcium imaging were presented with mechanical vibrations and light. (A, C) Percentage of excited (2 STD above baseline ) or inhibited (2 STD below baseline) Hb neurons for no injected fish (grey) or control-injected (cyan) fish in response to mechanical vibrations (A) or light (C) stimulation. (A) or light (C) stimulation. Note that there is no difference between the groups. No Injection n = 9, Control n = 7. (B, D) Average $\Delta$F/F amplitude (%) during the response period of all neurons in Hb per fish. No Injection n = 9, Control n = 7. (E) Percentage of unimodal Hb neurons that respond exclusively to either light or vibrations versus multimodal (M) neurons responding to both light and vibrations. Note that there is no difference between the groups. No Injection n = 9, Control n = 7. (F) Pearson’s correlation of multi-neuronal response vectors in the Hb for mechanical vibrations and light. Note that there is no difference between the groups. No Injection n = 9, Control n = 7. (G) Pairwise Pearson’s correlation of Hb neurons during spontaneous activity as a function of distance ($\mu$m) between each neuron pair in no injection fish (grey) versus control-injected fish (cyan). Note that CPPG-injected animals exhibit stronger correlations over longer distances. Shadow represents SEM. (*p < 0.05, ** p < 0.01, *** p < 0.001), tailed Wilcoxon rank sum test. Error bars present +/-SEM.

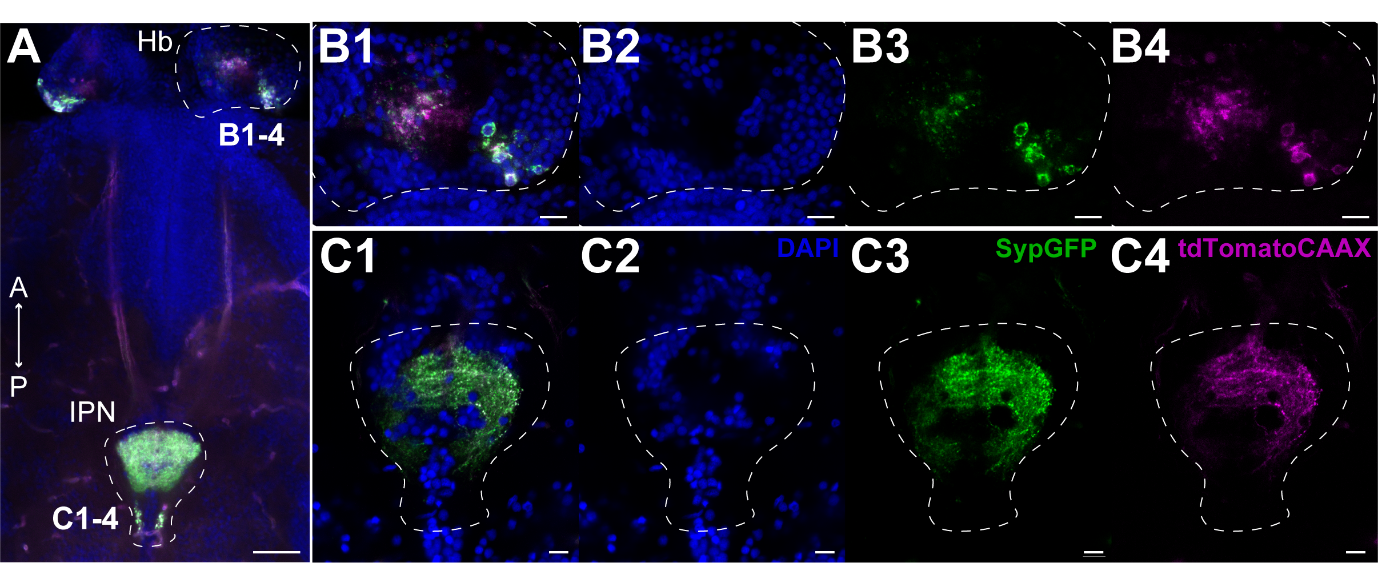

Supplementary Figure 3: Narp-population of Hb has presynapses both in IPN as well as Hb. Confocal images of *Tg(narp:GAL4VP16*;*UAS:Syp–GFP-T2A-tdTomato-CAAX )* juvenile zebrafish fixed, cleared and DAPI stained. Colors represent DAPI (blue), SypGFP (green) and tdTomatoCAAX (magenta). (A) Full image of Hb and IPN merged images of three channels. Dotted square indicated the Hb zoomed in H1. (B) Zoom into Hb (delineated with dotted line) from A. Merged (B1), DAPI (B2, blue), tdTomatoCAAX (B3, magenta), SypGFP (B4, green). Arrows are pointing at examples of individual presynapses coming from the Narp population staying inside the neuropil of Hb. (C) Zoom into IPN (delineated with dotted line) from A. Merged (C1), DAPI (C2, blue), tdTomatoCAAX (C3, magenta), SypGFP (BC4, green). Scale bar represents 50 (A) or 10 (B-C) $\mu$m.

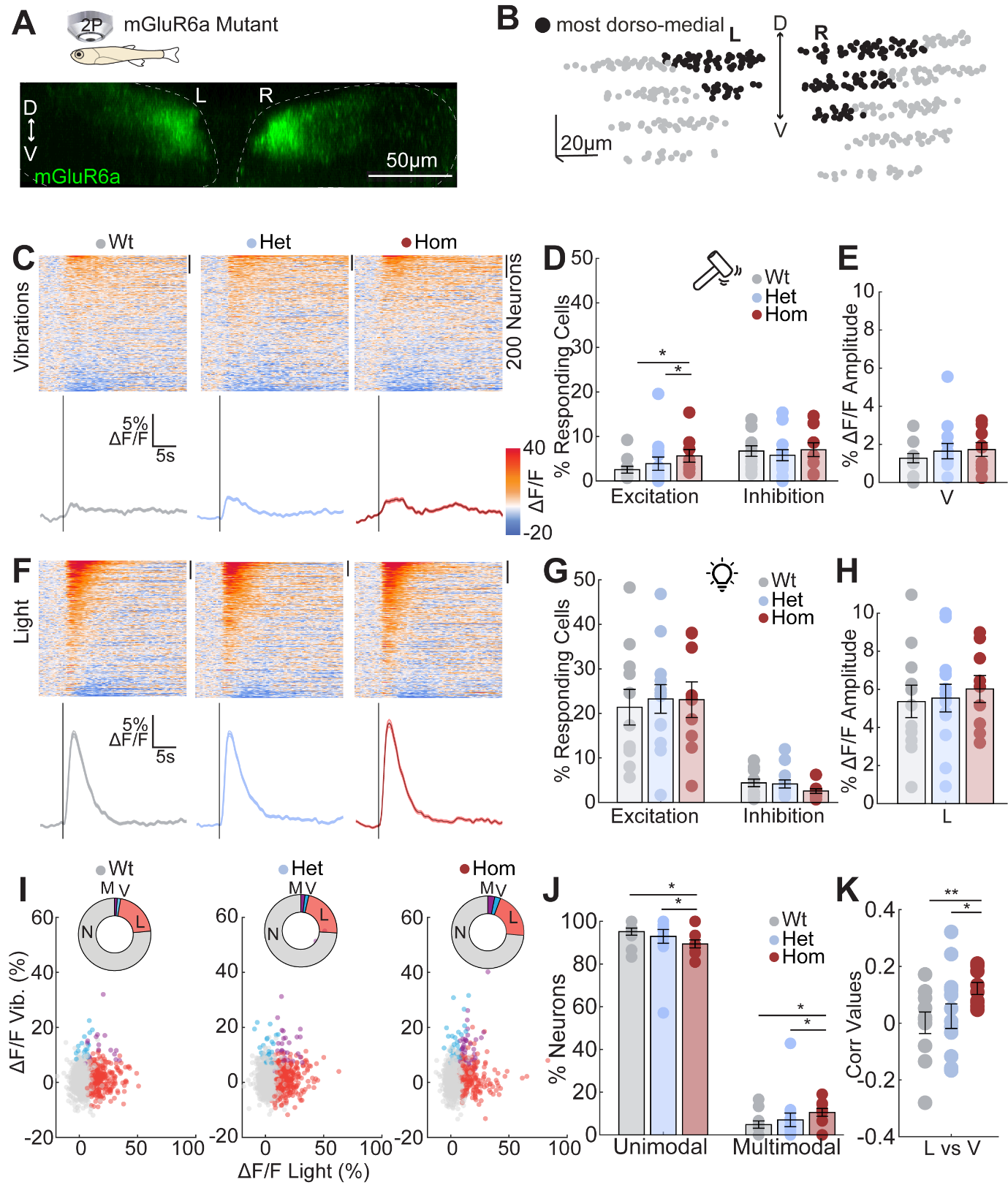

Supplementary Figure 4: Sensory responses and selectivity in Hb neurons exposed to mechanical vibrations and light stimulations in mGluR6a mutants. (A) Coronal confocal image of Hb that was stained by hybridization chain reaction for mGluR6a (green). The two Hb are delineated with dotted lines. Scale bar represents 50 $\mu$m. (B) 3D-representative example of a Wt fish where the most dorsomedial cells (black) show similar location as compared to the mGluR6a expression pattern seen in A. (C,F) Heatmaps represent the time courses of all Hb calcium signals ($\Delta$F/F) in responses to vibrations (C) or light (E) recorded. by two-photon calcium imaging in mGluR6a mutants expressing *Tg(elavl3:GCaMP6s* Left: Wt (n = 1454 neurons, 11 fish), middle: Het (n = 1726 neurons, n = 13 fish) or right: Hom (n = 1182 neurons, n = 9 fish) are sorted according to their mean activity in the response period (10s after stimulus onset). Warm colors indicate excitation, cold colors represent inhibition. Average traces of all Hb neurons are below each heatmap. Stimulus onset is indicated by a line. Shadow represents SEM. (D, G) Percentage of excited (2 STD above baseline) or inhibited (1 STD below baseline) Hb neurons Wt (grey), Het (blue) or Hom (red) fish in response to mechanical vibrations (D) or light (G) stimulation. Note that significantly more neurons in the Hom respond to vibrations compared to Wt and Het. Wt n = 11, Het n = 13, Hom n = 9 (E,H) Average $\Delta$F/F Amplitude (%) during the response period of all neurons in Hb per fish. (I) Responses of individual Hb neurons to mechanical vibration (blue), light (red) or both (magenta) for control and CPPG injected fish. Donut chart represents the ratio of Hb neurons and their response type (2 STD above baseline levels). N: non-responding, V: only vibrations, L: only light, M: both vibrations and light. (H) Percentage of unimodal Hb neurons that respond exclusively to either light (L) or vibrations (V) versus multimodal (M) neurons responding to both light and vibrations. Significantly less Hb neurons in the Hom are selective for one of the two stimulus modalities (unimodal), but instead more neurons are multimodal. (I) Pearson’s correlation of multi-neuronal response vectors in the Hb for mechanical vibrations and light. Sensory responses in the Hom are significantly more correlated and hence more similar to each other. (*p < 0.05, ** p < 0.01, *** p < 0.001), tailed Wilcoxon rank sum test. Error bars present +/-SEM.
